# Clustering Digestive Tract Tumors Using Transcriptomic and Mutation Data

**DOI:** 10.1101/2025.01.13.632722

**Authors:** Dwayne G. Tally, Polina Bombina, Jake Reed, Jeffrey Kinne, Lynne V. Abruzzo, Kevin R. Coombes, Zachary B. Abrams

**Affiliations:** Department of Informatics Indiana University Bloomington, IN 47408, USA; Department of Biostatistics, Data Science, and Epidemiology Georgia Cancer Center at Augusta University Augusta, GA 30912, USA; Department of Computer Science Indiana State University, Terre Haute, IN 47809, USA; Department of Pathology Medical University of South Carolina Charleston, SC 29425 USA; Institute for Informatics, Data Science & Biostatistics Washington University St. Louis, MO 63110, USA

## Abstract

Digestive tract cancers, like most other cancers, are usually categorized based on cell or tissue of origin. Molecular clustering based on the transcriptome often produces the same classification. We developed a new method, Newmanization, to reduce underlying tissue signals from transcriptomic analysis. To test our method, we downloaded data on 1635 samples of digestive tract cancers from The Cancer Genome Atlas. The available data includes transcriptomic data, by RNA-Seq, as well as binary mutation allele frequency data by whole exome sequencing. We compared, using silhouette widths and visualization by dimension reduction plots, the effectiveness of Newmanized transcriptome and mutation data to separate digestive tract cancers. The Newmanized transcriptome clusters have clearer separation and larger average silhouette widths. Feature analysis of each cluster for Newmanized transcriptomic data and mutation data revealed that clusters determined with Newmanized data contained more mRNAs present at higher frequencies than clusters defined by mutation data. This suggests that the Newmanized method holds great potential for advancing personalized transcriptomic medicine.

## 1 Introduction

Cluster analysis has been an integral part of the analysis of gene expression data since its earliest days [1]. In general, the goal of these analyses has been to discover subtypes of already known cancer types, usually characterized by the tissue or anatomic site of origin. Beginning in the mid-2000’s, as The Cancer Genome Atlas (TCGA) project accumulated data on cancers of different types, many of their initial publications applied clustering to find subtypes within individual cancer types, using either single- or multi-omic data sets. They typically reported roughly three to six subtypes of every cancer (ovarian, three or four [2]; colorectal, four [3]; breast, four [4]; lung squamous cell, four [5]; clear cell renal cell, four [6]; glioblastoma, six [7]).

The implicit assumption underlying this class of applications of clustering to gene expression data is that different types of cancer are each characterized by a small set of “drivers” comprised of genomic (somatic mutations and copy number changes) and epigenetic lesions. These lesions then impose a dominant “signal” on the gene expression patterns that reflect the specific ways that some critical pathways and hallmarks of cancer are broken in those cancer types. The mutations, in particular, have precipitated the growth of “personalized medicine” in cancer by providing therapeutic targets that are detectable in individual tumors.

Once TCGA grew large enough to contain data on cancer types from many different organs, researchers began applying clustering as part of their “pan-cancer” analyses [8]. If that implicit assumption were correct, one might expect the pan-cancer clusters to cross the boundaries between organs, since (a) the gene expression in each cancer should reflect the mutation patterns and (b) many of the same cancer driver genes are mutated in different kinds of cancer. Instead, it soon became clear that the “type of cancer” signal was unlikely to be the strongest part of the gene expression profile.

For example, Hoadley and colleagues [9] reported that: “We performed molecular clustering using data on chromosome-arm-level aneuploidy, DNA hypermethylation, mRNA, and miRNA expression levels and reverse-phase protein arrays, of which all, except for aneuploidy, revealed clustering primarily organized by histology, tissue type, or anatomic origin.” Wu and colleagues [10] found that “the pan-cancer analysis results show that the cancers of different tissue origins are generally grouped as independent clusters, except squamous-like carcinomas.” Taskesen and colleagues drew similar conclusions from their multi-omic clustering study [11]. The same result has been found in single omic studies, including methylation profiles [12], mutational signatures [13], and in proteomic patterns from The Cancer Proteomic Atlas (TCPA) [14].

The dominance of the signal from the cell or tissue of origin persisted even when one restricted to the expression of smaller sets of genes. We previously performed a clustering analysis across TCGA cancer types by using the gene expression patterns of a representative subset of transcription factors. We found that the primary driver of cluster formation was the cell or tissue of origin [15]. We obtained similar results when clustering samples based on the expression of a representative set of microRNAs [16]. Careful perusal of the t-SNE plots in those papers (and in many of the papers cited above) suggests that it is the cell, rather than the tissue, of origin that dominates. For example, non-small-cell lung cancer has long been separated into two histological subtypes: adenocarcinoma or squamous cell carcinoma. These subtypes were treated as different primary types by TCGA, and they cluster separately in pan-cancer analyses. Moreover, the squamous cell lung cancers tend to cluster relatively close to, but not overlapping, squamous cell cancers from other sites, including head-and-neck or cervix. Esophageal cancer also has two histological subtypes. Esophageal adenocarcinomas tend to cluster with stomach adenocarcinoma, and esophageal squamous cell carcinomas tend to cluster with head-and-neck squamous cell tumors [17].

One way to get around the dominance of the cell-of-origin signal can be found in a report from Akbani and colleagues, who clustered TCGA samples using reverse-phase protein array (RPPA) data [18]. Tissue of origin was found to be a strong driver of clustering when the data were processed using standard normalization. However, this effect was significantly reduced after applying tissue-specific median centering to remove the signal from tissue types. A potential problem with this approach, however, is that we have no idea exactly what is being thrown away along with the tissue-of-origin signal. Important properties that are common to the entire set of cancers may also be discarded.

In this paper, we introduce the “Newman banked statistic”, which we compute using a “bank” of normal control samples based on the tissue of origin for each type of cancer. We standardize the gene expression patterns in tumor samples using the observed mean and standard deviation of the normal bank. Using this method, we have control over our modifications to the full transcriptomic profile, ensuring that we only remove the signal corresponding to average normal tissue expression. The resulting data for each gene is then measured in how many standard deviation units it is away from the normal mean. By setting a threshold in such units, we can convert the standardized expression data to binary data, with “0” representing “in the normal range” and “1” representing “outside the normal range”. We refer to the entire process of standardization and dichotomization as “Newmanization”. We suspect that Newmanized transcriptomic data may give a clearer picture of the functional effects of important drivers on gene expression by automatically filtering out the weaker effects of passenger mutations.

Our goal here is to test two hypotheses. First, we hypothesize that clustering using the binary mutation data should produce clusters based on mutation patterns rather than cell of origin. Second, we hypothesize that clustering using the binary Newmanized transcriptomic data will produce clearer and more interpretable clusters. To test these hypotheses, we will use six cancers from anatomic sites along the digestive tract (head-and-neck, esophagus, stomach, pancreas, colon, and rectum) from the collection of TCGA data. As with most cancers, digestive tract cancers are usually categorized based on cell type and location [19]. However, colon and rectal cancers are almost indistinguishable based on their usual transcriptomic profiles [3,15]. These relationships may reflect only the normal expression patterns in the cell of origin. Our hypotheses are that mutation patterns, especially as reflected in Newmanized transcriptomic data, will perform better at identifying meaningful molecular subtypes that cut across the boundaries defined by anatomical sites. Digestive tract cancers provide an ideal context to test this approach, given their diversity in tissue types and mutation patterns. This diversity should reveal whether unsupervised mutation clustering or Newmanized clustering can uncover meaningful molecular subtypes independent of cell of origin. If successful, this study could pave the way for advances in precision oncology by offering insights into cross-tissue cancer subtypes and identifying novel therapeutic targets.

## 2 Methods

### 2.1 Data

We obtained transcriptomic and mutation allele frequency data from TCGA. The transcriptomic data contains diseases associated with colon (COAD), rectum (READ), pancreas (PAAD), stomach (STAD), esophagus (ESCA), and head and neck (HNSC). **Table 1** shows the number of samples per digestive tract disease type. The transcriptomic data contains continuous measurements of 20531 features (i.e., genes) on samples from 1635 patients. The mutation data set is binary (wild-type or mutated) for the same 1635 patients but only has 563 features (i.e., potentially mutated genes).

**Table 1:**
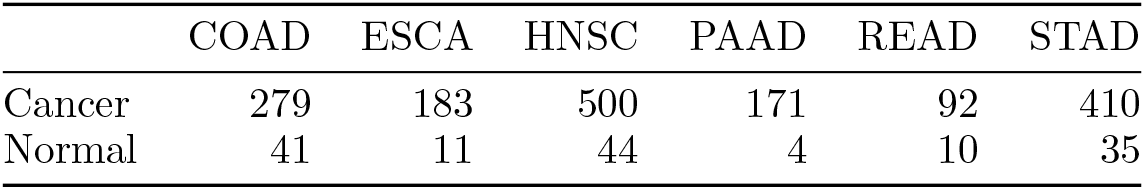
Number of samples by digestive tract cancer site of origin.

|  | COAD | ESCA | HNSC | PAAD | READ | STAD |
| --- | --- | --- | --- | --- | --- | --- |
| Cancer | 279 | 183 | 500 | 171 | 92 | 410 |
| Normal | 41 | 11 | 44 | 4 | 10 | 35 |

### 2.2 Software

All analyses were performed using version 4.1.1 of the R statistical software environment. We used a variety of public R packages: NewmanOmics version 1.0.12, Mercator version 1.1.5 [20], and Polychrome version 1.5.1 [21]. Clustering was performed using the implementation of hierarchical clustering with Ward’s linkage rule in the hclust function in the stats package from base R. Visualization was performed using the Mercator interface to the t-SNE algorithm in version 0.17 of the Rtsne R package [22]. All R packages are available from the Comprehensive R Archive Network (CRAN) [23].

### 2.3 Newman Bank Test

We used the NewmanOmics package to process the TCGA transcriptomic data. The underlying logic of the Newman bank statistic is derived from D. Newman’s 1939 studentized range statistic [24]. Since Newman’s statistic requires an independent estimate of the standard deviation, we converted the expression of gene *g* by standardizing it using a cohort (or “bank”) of normal control samples:

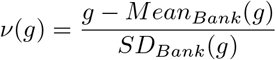

After this procedure, the expression of each gene in each tumor sample is recorded as the number of (normal control) standard deviations it lies away from the (normal control) mean. As a result, we obtain detailed information on each individual patient about the set of genes whose expression is abnormal.

### 2.4 Dichotomization and Clustering

After standardizing the usual transcriptomic data, we converted the resulting matrix into binary, based on a cutoff value measured in standard deviations. Specifically, any gene whose expression was within three standard deviations of the mean was coded as “0”, and any gene whose expression was more than three standard deviations above or below the mean was coded as “1”.

We then applied the Mercator package to both the Newmanized transcriptome data and the mutation data. Since both data sets are binary, we used the Jaccard distance metric [25]. The Jaccard similarity, *S*_*J*_, between two binary vectors is defined as

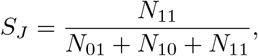

where *N*_*ij*_ counts the number of times a value equals *i* in the first vector and *j* in the second vector. Then the Jaccard distance is defined as *D*_*J*_ = 1 − *S*_*J*_. One advantage of using the Jaccard distance instead of other measures of difference between binary vectors is that it ignores the non-informative “0-0” matches when events (like mutations) are relatively rare and/or their absence is less important than their presence.

We applied hierarchical clustering to select six clusters from the Jaccard distance matrices, one cluster for each known disease category. We used silhouette widths to measure the intrinsic quality of clustering [26,27]. Values of the silhouette width lie between −1 and +1. When the silhouette width is positive, then the clustering is good. When the score is negative, the clustering is poor. We used Polychrome to generate standard colors in order to display the clusters in a t-SNE plot [21,22].

### 2.5 Interpretation

We used summary statistics to estimate the frequency of each event (mutated gene or abnormally expressed gene) in each cluster. In order to identify genes that were substantially different between clusters, we used both gene-by-gene chi-squared tests on the binary Newmanized data and gene-by-gene analysis of variance (ANOVA) on the continuous *ν*-values.. Finally, we performed gene enrichment analysis using ToppGene (https://toppgene.cchmc.org/) [28].

## 3 Results

### 3.1 Comparing Clusters From Usual and Newmanized Transcriptomic Data

As shown in Figure 1A, the usual transcriptomic clusters are defined mainly by the cell type or tissue of origin. There are essentially four clusters, consisting of pancreas (PAAD), colorectal (COAD, READ), stomach (STAD), and head-and-neck (HNSC). The esophageal tumors (ESCA) cluster either with stomach (if adenocarcinomas) or head and neck (if squamous cell cancers). However, clusters learned from the binary Newmanized transcriptome data are not defined by tissue (Figure 1B), though they show a similar degree of separation to the usual transcriptomic clusters. In this plot, stomach tumors are mostly clustered apart from all other DT cancers. All the other diseases are split between at least two clusters, and each of the four additional clusters combines multiple tumor types. One cluster contains esophagus and head and neck; a second contains esophagus and colon; a third contains head and neck and pancreas; and the fourth contains pancreas and rectal.

**Figure 1:**
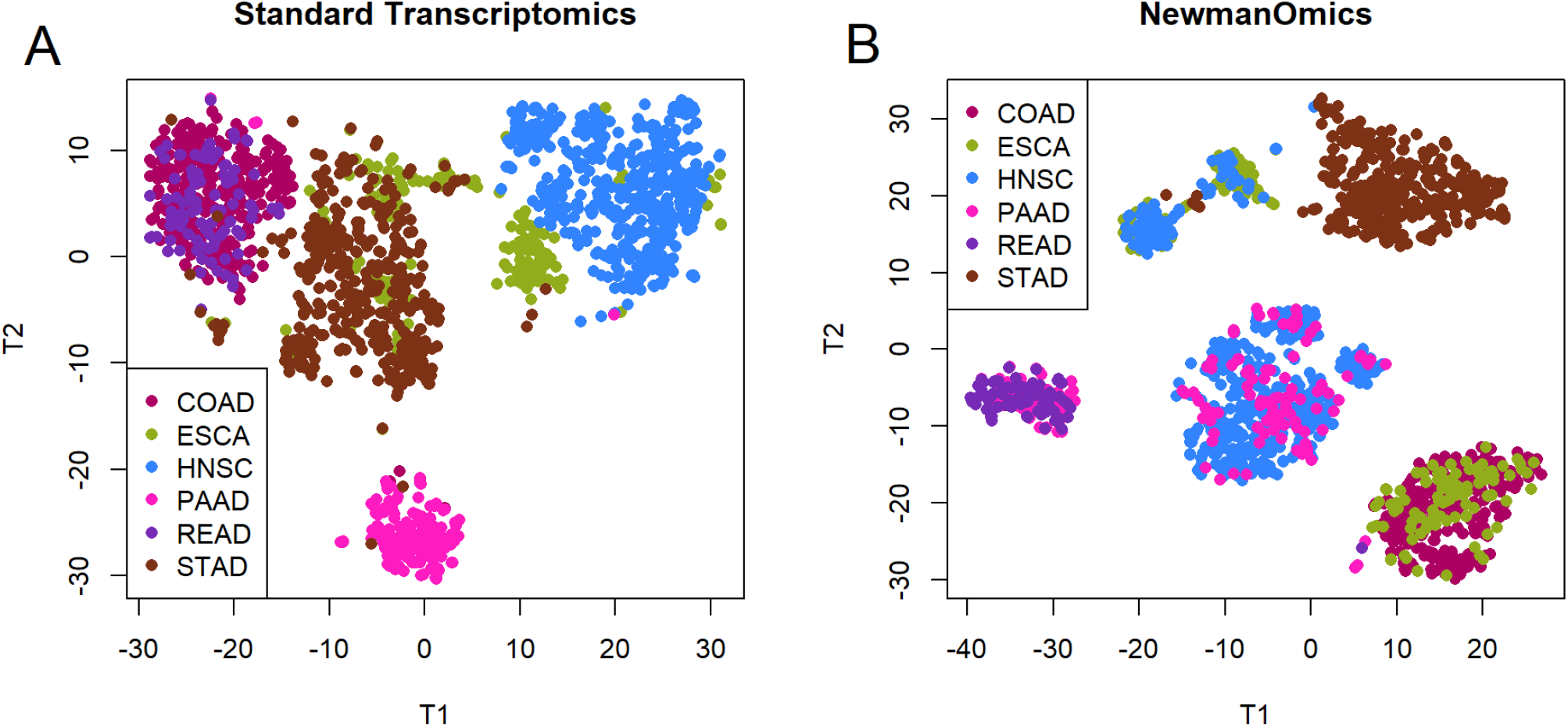
t-SNE comparison between (A) standard and (B) Newman transcriptomic data. Colors relect the tissue or cell of origin.

### 3.2 Comparing Newman Transcriptomic and Mutation Allele Frequency

The next figure (Figure 2) is a four-part panel that uses t-SNE to visualize hierarchical clusters and tissue types between Newman transcriptomic and mutation allele frequency clustering. The t-SNE plots in Figures 2A and 2C are the same underlying plots derived from the Newmanized transcriptome data. Figure 2A colors the points based on unsupervised hierarchical clustering, while Figure 2C uses colors to indicate the tissue of origin and is identical (by intention) to Figure 1A. The six clusters in Figure 1A match one’s expectations from the structure of the t-SNE plot. Meanwhile, Figures 2B and 2D show the corresponding plots based on the binary mutation data. The boundaries between clusters in Figure 2B are quite blurry, with very little in the form of recognizable, separable clusters. Figure 2D uses different colors to show the tissue of origin of the tumor samples. There is some indication of overlap between hierarchical clusters and tissue type, but it is not as strong as that seen with the transcriptomic data in Figure 1A.

### 3.3 Silhouette Width

Figure 3 visualizes the silhouette width plots for hierarchical clusters using either (A) Newmanized transcriptomic or (B) mutation data. The silhouette width of the Newmanized data visually shows a higher rate compared to the silhouette width of the mutation data The mean silhouette width of the Newmanized transcriptomic data is substantially higher (*μ* = 0.0828) than the mean silhouette width of the mutation data (*μ* = 0.0039). The plot for Newmanized transcriptomic data shows that almost every sample was categorized into the correct cluster, except for the fourth (magenta) cluster where some samples had negative values. Mutation silhouette widths are mostly negative except for the second (dark olive), fourth (purple) and fifth (lavender) clusters. This indicates that almost every sample in the other clusters could have been categorized better.

### 3.4 Common Features Associated with Hierarchical Clusters

Table 2 lists the top five features most frequently abnormally expessed or mutated in each cluster from each of the binary data sets. Panel A (columns 1-3) lists mRNA features when clusters are found from Newmanized transcriptomic data. Panel B (columns 4-6) lists mRNA features when clusters are created from mutation data. Panel C (columns 7-9) lists mutated genes when clusters are found from Newmanized data. Panel D (columns 10-12) lists mutated genes when clusters are created from mutation data. Features listed in Panel A are present at a much higher percentage than those in any other panel.

Abbreviations: RN = mRNA features associated with clusters defined by Nemmanomics. RM = mRNA features associated with clusters defined by mutations. MN = Mutated genes associated with clusters defined by Nemmanomics. MM = Mutated genes associated with clusters defined by mutations.

**Figure 2:**
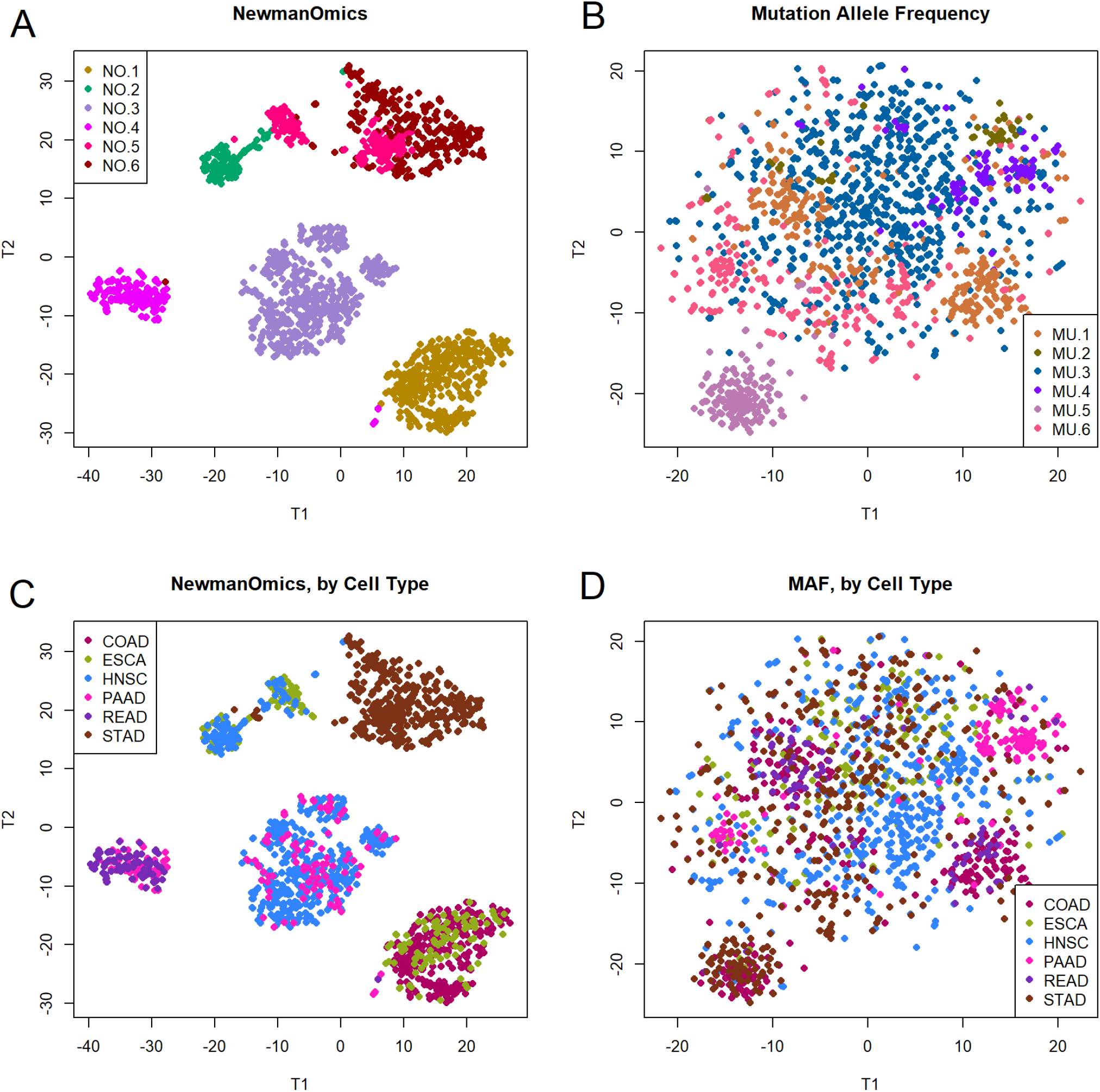
t-SNE graphical analysis of binary Newmanized transcriptomic (A,C) and mutation allele frequency (B,D) data based on hierarchical clusters (A,B) or colored by disease type (C,D). Both (A) and (B) are colored by hierarchical clusters and (C) and (D) are colored by known diseases. (A) shows Newmanized transcriptomic data. (B) shows mutation allele frequency hierarchical clusters. (C) shows the transcriptomic clusters but colored by the disease category. (D) shows the mutation allele frequency clusters but colored by disease type.

**Figure 3:**
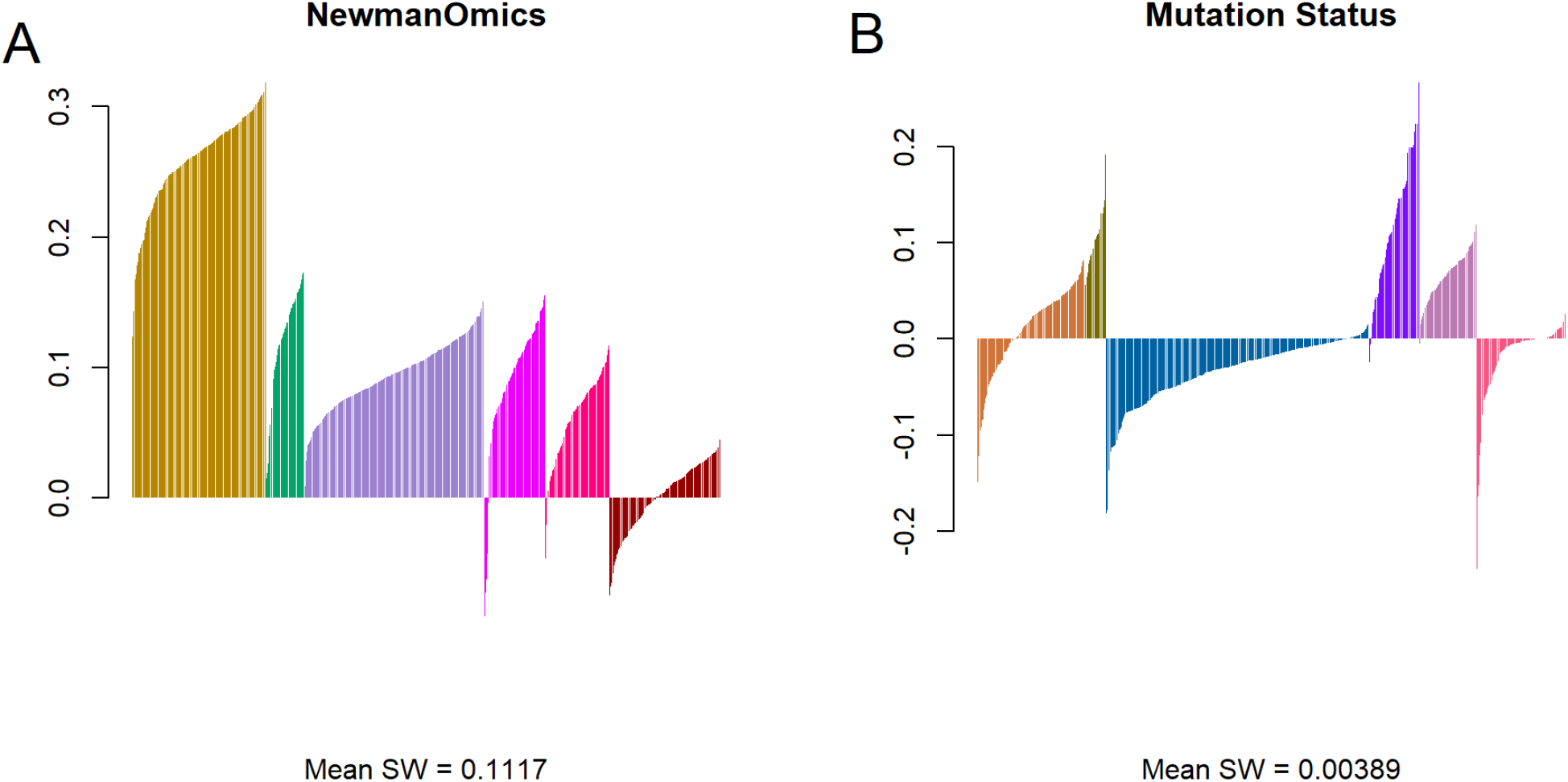
Silhouette widths plots for hierarchical clustering of (A) Newmanized transcriptomic and (B) mutation allele frequency data.

**Table 2:** Top five significant genes, by cluster.

| RN.# | RN.Gene | RN.% | RM.# | RM.Gene | RM.% | MN.# | MN.Gene | MN.% | MM.# | MM.Gene | MM.% |
| --- | --- | --- | --- | --- | --- | --- | --- | --- | --- | --- | --- |
| 1 | CA7 | 100.0 | 1 | XIST | 86.2 | 1 | TP53 | 61.6 | 1 | APC | 74.5 |
| 1 | OTOP2 | 99.7 | 1 | ADH1B | 82.2 | 1 | APC | 56.2 | 1 | TP53 | 67.8 |
| 1 | KRT80 | 99.7 | 1 | SFRP1 | 80.5 | 1 | KRAS | 33.3 | 1 | KRAS | 45.0 |
| 1 | GCG | 99.7 | 1 | COL11A1 | 79.2 | 1 | PIK3CA | 23.9 | 1 | PIK3CA | 26.8 |
| 1 | BEST4 | 99.7 | 1 | UGT1A10 | 78.2 | 1 | FAT4 | 21.2 | 1 | FAT4 | 22.5 |
| 2 | KRT5 | 100.0 | 2 | MUC5B | 83.3 | 2 | TP53 | 76.9 | 2 | SMAD4 | 98.3 |
| 2 | TRIM29 | 99.1 | 2 | DMBT1 | 73.3 | 2 | LRP1B | 20.4 | 2 | TP53 | 65.0 |
| 2 | GKN1 | 99.1 | 2 | MMP1 | 71.7 | 2 | CDKN2A | 19.4 | 2 | KRAS | 48.3 |
| 2 | ATP4B | 99.1 | 2 | KRT7 | 71.7 | 2 | KMT2D | 16.7 | 2 | APC | 16.7 |
| 2 | ATP4A | 99.1 | 2 | CEACAM6 | 71.7 | 2 | NOTCH1 | 14.8 | 2 | RNF43 | 10.0 |
| 3 | CRISP3 | 95.6 | 3 | ADH1B | 83.8 | 3 | TP53 | 66.0 | 3 | TP53 | 77.8 |
| 3 | PRH2 | 94.2 | 3 | MYOC | 83.5 | 3 | FAT1 | 19.0 | 3 | LRP1B | 20.2 |
| 3 | SMR3B | 93.4 | 3 | PLIN4 | 83.1 | 3 | CDKN2A | 18.4 | 3 | FAT1 | 12.9 |
| 3 | PRR4 | 92.4 | 3 | MAL | 82.6 | 3 | PIK3CA | 15.4 | 3 | KMT2D | 11.0 |
| 3 | MUC7 | 92.4 | 3 | PI16 | 79.7 | 3 | KRAS | 13.8 | 3 | CDKN2A | 9.7 |
| 4 | XIST | 97.0 | 4 | MUC5B | 89.4 | 4 | TP53 | 71.6 | 4 | TP53 | 90.8 |
| 4 | AGR2 | 96.4 | 4 | PIGR | 79.4 | 4 | KRAS | 55.0 | 4 | KRAS | 58.2 |
| 4 | CEACAM6 | 94.7 | 4 | ADH1B | 79.4 | 4 | APC | 46.7 | 4 | CDKN2A | 44.0 |
| 4 | TMPRSS4 | 94.1 | 4 | DMBT1 | 75.9 | 4 | SMAD4 | 17.8 | 4 | RNF213 | 7.8 |
| 4 | CTSE | 92.3 | 4 | PLIN4 | 73.0 | 4 | FAT4 | 12.4 | 4 | SMAD4 | 6.4 |
| 5 | MYOC | 96.6 | 5 | CLEC3B | 93.7 | 5 | TP53 | 59.6 | 5 | KMT2D | 64.8 |
| 5 | PLP1 | 96.1 | 5 | MYOC | 93.1 | 5 | LRP1B | 32.0 | 5 | ARID1A | 62.3 |
| 5 | EPCAM | 96.1 | 5 | SCNN1B | 91.2 | 5 | ARID1A | 26.4 | 5 | FAT4 | 57.9 |
| 5 | PRIMA1 | 95.5 | 5 | SCARA5 | 89.9 | 5 | KMT2D | 22.5 | 5 | RNF213 | 50.3 |
| 5 | NCAM1 | 95.5 | 5 | PI16 | 89.9 | 5 | FAT4 | 21.9 | 5 | LRP1B | 49.1 |
| 6 | MAL | 93.5 | 6 | MAL | 84.3 | 6 | TP53 | 47.7 | 6 | PIK3CA | 22.6 |
| 6 | COL10A1 | 91.2 | 6 | COL10A1 | 77.0 | 6 | LRP1B | 23.4 | 6 | ARID1A | 16.5 |
| 6 | ESM1 | 88.3 | 6 | PLIN4 | 75.4 | 6 | ARID1A | 19.5 | 6 | CASP8 | 11.3 |
| 6 | CST1 | 88.3 | 6 | DMBT1 | 74.6 | 6 | FAT4 | 16.6 | 6 | TP53 | 10.9 |
| 6 | PGA4 | 87.7 | 6 | MYOC | 73.8 | 6 | PIK3CA | 13.0 | 6 | FAT4 | 10.5 |

#### 3.4.1 Chi-Squared Tests

Mutated gene features in Panel D (clustered by mutation) of Table 2 are typically at higher frequencies than those in Panel C (clustered by Newmanized transcriptome). While it is not terribly surprising that the most frequently mutated genes occur at higher rates when the clusters are defined by mutation patterns, one must take special care when trying to interpret these results. For example, *TP53* is the most commonly mutated gene across the entire combined data set, where it is mutated in 62.1% of all samples. It is listed as one of the five most commonly mutated genes in Cluster 6 of the mutation hierarchical data, but it is only present in 10.9% of those samples. This observation suggests that *TP53* mutations are more notable for their absence in these samples than their presence. To address this issue, we performed chi-squared tests for each gene in each cluster to determine which gene mutations are present at higher rates than would be predicted by their overall mutation rate.

Mutated genes associated with NewmanOmics clusters:

- NO.1 (279 COAD, 91 ESCA cases): ***APC*** (Overall: 22.1%; Here: 56.5%, *χ*^2^ = 373.9), ***BRAF*** (Overall: 5.0%; Here: 12.7%; *χ*^2^ = 64.1), ***FAT4*** (Overall: 14.4%; Here: 21.4%; *χ*^2^ = 50.6), plus ten others
- NO.2 (91 READ, 68 PAAD cases): ***KRAS*** (Overall: 19.8%; Here: 54.1%; *χ*^2^ = 193.5), ***NRAS*** (Overall: 1.5%; Here: 6.3%; *χ*^2^ = 36.4)
- NO.3 (404 STAD cases): ***ARID1A*** (Overall: 11.4%; Here: 25.8%; *χ*^2^ = 115.0), ***CDH1*** (Overall: 3.4%; Here: 9.6%; *χ*^2^ = 64.4), ***ACVR2A*** (Overall: 5.0%; Here: 10.8%; *χ*^2^ = 63.2), plus ten others
- NO.4 (173 HNSC, 54 PAAD cases): ***CDKN2A*** (Overall: 9.8%; Here: 19.7%; *χ*^2^ = 83.5), ***HRAS*** (Overall: 2.0%; Here: 5.6%; *χ*^2^ = 35.7), ***FAT1*** (Overall: 11.9%; Here: 19.7%; *χ*^2^ = 30.4) ***CASP8*** (Overall: 5.0%; Here: 10.2%; *χ*^2^ = 29.4)
- NO.5 (91 HNSC, 90 ESCA cases): ***NFE2L2*** (Overall: 3.3%; Here: 8.0%; *χ*^2^ = 30.6), ***TP53*** (Overall: 62.1%; Here: 79.1%; *χ*^2^ = 29.3)
- NO.6 (235 HNSC, 49 PAAD cases): ***NOTCH1*** (Overall: 9.1%; Here: 15.3%; *χ*^2^ = 29.1)

Mutated genes associated with mutation clusters:

- MU.1 (150 COAD, 62 READ, 76 other cases): ***APC*** (Overall: 22.1%; Here: 74.5%; *χ*^2^ = 498.8)
- MU.2 (27 PAAD; 12 STAD; 21 other cases): ***SMAD4*** (Overall: 8.9%; Here: 98.3%; *χ*^2^ = 579.5)
- MU.3 (318 HNSC, 184 STAD, 136 ESCA, 91 other cases): ***TP53*** (Overall: 62.1%; Here: 77.8%; *χ*^2^ = 175.0)
- MU.4 (74 PAAD, 42 HNSC, 25 other cases): ***KRAS*** (Overall: 19.8%; Here: 58.2%; *χ*^2^ = 350.1), ***CDKN2A*** (Overall: 9.8%; Here: 44.0%; *χ*^2^ = 200.5)
- MU.5 (81 STAD, 56 COAD, 22 other cases): ***ACVR2A*** (Overall: 5.0%; Here: 40.3%; *χ*^2^ = 438.8), ***ZFHX3*** (Overall: 6.2%; Here: 45.3%; *χ*^2^ = 436.1) ***ARID1A*** (Overall: 11.4%; Here: 62.3%; *χ*^2^ = 428.6) plus more than a hundred others
- MU.6 (90 HNSC, 88 STAD, 70 other cases): None.

The large number of genes associated with cluster MU.5 suggests that those samples may simply have more mutations than other cases. To test this idea, we plotted the distribution of the number of mutated genes per sample within clusters (Figure 4). Most samples have fewer than ten mutated genes, but the samples in cluster MU.5 have closer to 100 mutations each. These samples appear to be distributed across Newmanized clusters NO.1 and NO.3, both of which have more associated mutated genes than other Newmanized clusters. We also note that there are 25 samples that have no mutated genes; all 25 are assigned to cluster MU.6 (which has no significantly associated genes). In the Newmanized clusters, these samples are scattered loosely across all six clusters.

### 3.5 Gene Enrichment

In order to understand the expression differences between the six clusters using the Newmanized transcriptome data, we performed two analyses. First, we performed gene-by-gene ANOVA on the continuous *ν*-values. Second, we performed gene-by-gene chi-squared tests on the binary values. The complete results of both analyses are presented in an Excel spreadsheet (**Supplemental Table 1**). In both cases, the false discovery rate (FDR) associated with nominal p-value cutoffs *p* < 0.01 satisfied FDR < 0.01. In order to select genes strongly associated with individual clusters, we restricted to the set of genes for which *p* < 0.01 for both the ANOVA F-test and the chi-squared test, and which were found to be abnormally expressed (i.e., outside the normal range) in at least 10% of all cases. This produced a list of 4677 genes. Each gene was allocated to the cluster that contributed the largest percentage of its cases to the set with abnormal expression. (By cluster, this procedure yielded 2719 genes for NO.1, 352 for NO.2, 104 for NO.3, 353 for NO.4, 652 for NO.5, and 505 for NO.6.) These gene sets were uploaded to the ToppGene web site for gene enrichment analysis, with parameters set to return at most 40 annotations per major ToppGene category. A full set of the enrichment tables produced by the web site can be found in an Excel spreadsheet (**Supplemental Table 2**).

**Figure 4:**
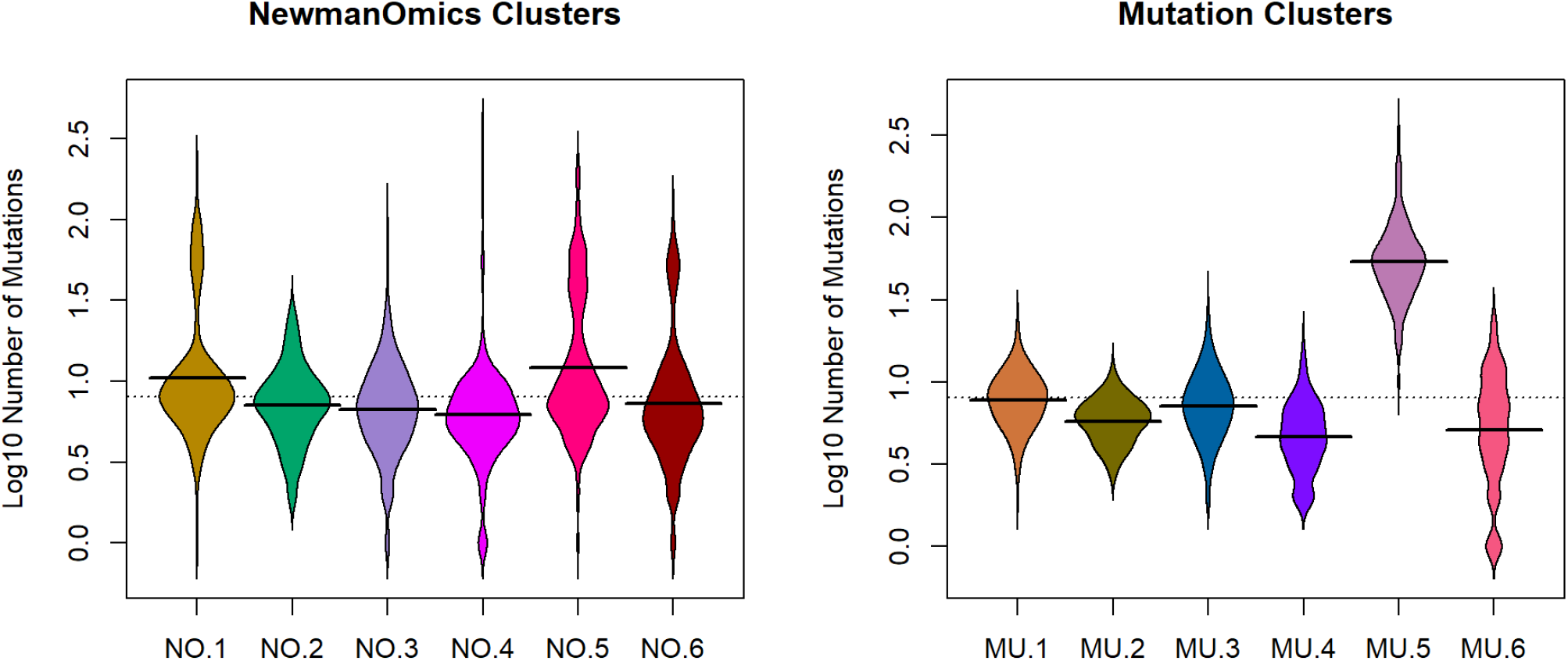
Mutation burden in clusters defined by (A) Newman transcriptomic data and (B) mutation data.

For visualization, we restricted to the five most significant annotations per cluster, along with any annotation that was associated with more than one cluster. The results for enriched pathways are shown in **Figure 5**. We note that multiple clusters are associated with the extracellular matrix, along with collagens, integrins, and adhesion.

#### 3.5.1 Differences Between Colon and Rectal Cancer

Next, in order to better understand the differences between cluster NO.1 (COAD and ESCA) and cluster NO.2 (READ and PAAD), we selected all genes significantly associated with either of those clusters for which they also had either the highest or lowest average expression of *ν*-values. We used this set of 1159 genes for gene enrichment analysis at ToppGene. Here the roles of the extracellular matrix, adhesion, collagens, and integrins were even more prominent. Among the pathway associations, 30 of the top 40 and 19 of the top 20 associated annotations were from this set of pathways. The same was true of 4 of the top 6 gene ontology molecular function annotations, 4 of the top 8 for biological process, and 6 of the top 10 for cellular component.

#### 3.5.2 Differences Between Subtypes of Esophageal Cancer

In NewmanOmics clustering, the esophageal cancers are split across two clusters, NO.1 (COAD and ESCA) and NO.5 (HNSC and ESCA). We compared this split to the known split into squamous cell carcinoma and adenocarcinoma (Table 3).

**Figure 5:**
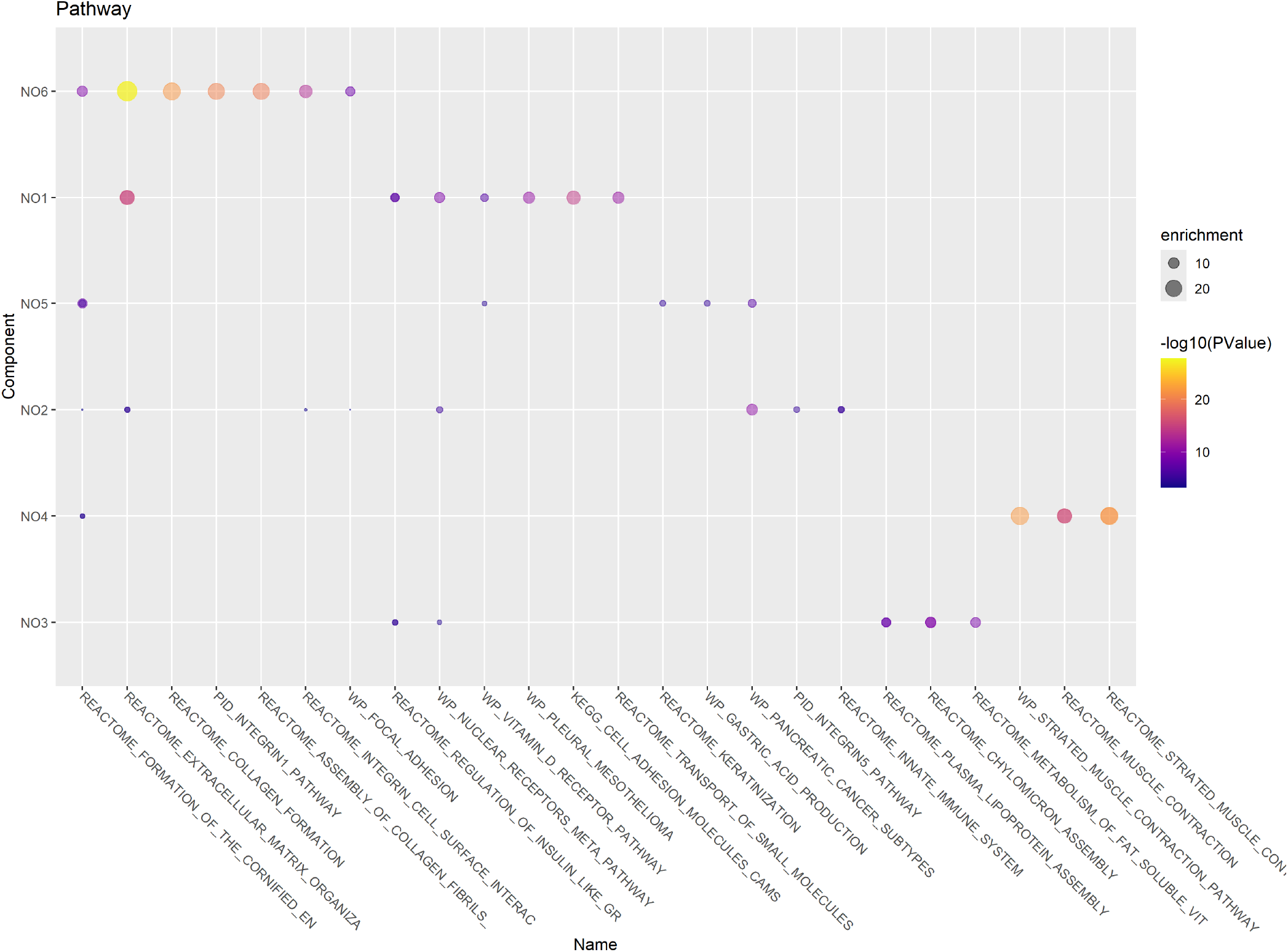
Summary of gene enrichment analysis for pathways, by Newmanomics clusters.

**Table 3:** Comparison between NewmanOmics clusters and histological cell types of esophageal cancer.

|  | NO.1 | NO.5 |
| --- | --- | --- |
| Adenocarcinoma | 57 | 6 |
| Squamous Cell Carcinoma | 29 | 28 |

While there is a statistically significant association of adenocarcinoma with Newmanized cluster NO.1 (*χ*^2^ = 22.344, *p* = 2.28 × 10^−9^), this association is far from perfect. To better understand which features are driving the Newmanized clustering, we performed a similar gene enrichment analysis comparing these NO.1 and NO.5 using 376 genes. We found significant differences that could be linked to two specific chromosomal cytobands. First, nine histones and nine HLA Class II genes in the HLA region on cytoband 6p21.3 were expressed at higher levels in cluster NO.5 compared to cluster NO.1. There were also associations with autoimmunity, T cell expression and receptors, and CTLA expression. Second, three APOBEC3 genes in cytoband 22q13.1 were also expressed at higher levels in cluster NO.5.

## 4 Discussion

Our first finding (Figure 1A) confirms previously published results that suggest that clustering and t-SNE plots of pan-cancer analyses are primarily driven by cell of origin [3,10,15,16]. This finding includes the observations that (a) colon cancer and rectal cancer are indistinguishable based on the usual transcriptomic profiles and (b) esophageal cancers split into two groups, one of which clusters with stomach adenocarcinomas and one of which clusters with head and neck squamous cell cancers. These exceptional cases are consistent with the known biological relations between the tissue types [17,29,30]. These findings also suggest that gene expression patterns that were already established in those cells before they became cancerous persisted after transformation. Moreover, those existing expression patterns appear to be the strongest signal in the data, dominating expression signatures arising from specific gene mutations in specific tumors.

Our second finding (Figures 2B, 2D, and 3B) is that using Jaccard distance metrics computed from binary mutation data does not lead to a clear clustering result. There are few clear boundaries, and very little in the way of well separated clusters. The silhouette width plot (Figure 3B), in particular, shows that about half the samples are not well clustered and may be more similar to other clusters than the one to which they are assigned.

In spite of the apparently low quality of many of the mutation clustering assignments, a deeper look at the genes associated to the clusters suggests that using the mutation data for clustering enables the discovery of patterns that cannot easily be found by any other unsupervised method. For example, the samples in cluster MU.5 have many more mutated genes on average than any other cluster. Note that MU.5 contains the lavender samples in the lower left corner of Figure 2B, which is the cluster most clearly separated from everything else. This observation is consistent with the silhouette width plot in Figure 3B, where all samples in MU.5 appear to be well clustered into the proper group. The cancer types contributing to MU.5 include 81 STAD, 56 COAD, 13 HNSC, and fewer than five samples from any of the other three cohorts. Higher mutation rates are often associated with microsatellite instability (MSI) [31–33], a phenomenon that is more common in stomach cancer and colon cancer than in other digestive tract cancer types [34,35]. Thus, clustering based on the binary mutation patterns may have detected a set of MSI cancer samples.

At the other extreme, mutation cluster MU.6 has no strongly associated mutated genes and, in fact, includes 25 cases that have no mutated genes. It is, therefore, not surprising that almost all samples in MU.6 are marked as poorly clustered in Figure 3B. All four of the remaining clusters are strongly associated with at least one or two mutated genes:

- The *APC* gene is mutated in almost 75% of the cases in MU.1 (where 70% of the cases are either COAD or READ). *APC* has long been known as one of the most frequently mutated genes in colorectal cancer [36].
- The *SMAD4* gene is mutated in more than 98% of the cases in MU.2. *SMAD4* is known to be a common mutation in pancreatic cancer [37,38], colorectal cancer [39,40], and gastric cancer [35].
- The *TP53* gene, which is known to be the most commonly mutated gene across cancer types [41], is mutated in almost 78% of the cases in MU.3 and 91% of the cases in MU.4. Both of these rates are higher than the overall level of about 62% of mutated cases in this data set.
- By the chi-squared test, MU.4 is also strongly associated with two other mutated genes, *KRAS* and *CDKN2A. KRAS* mutations are known to be relatively common in pancreatic cancer and colorectal cancer [42,43]. *CDKN2A* is known to be mutated in both pancreatic cancer [44] and head and neck cancer [45]. The co-occurrence of these two mutations in pancreatic cancer has also been reported previously [44].

It is not clear why clusters are so poorly separated when using the binary mutation data. Our first thought was that it might be a consequence of the presence of “passenger” mutations that blur the boundaries. However, the fact that the highly mutated MSI cases define the most clearly identifiable cluster may contradict that notion. It also appears that there are overlaps between the relatively small sets of most frequently mutated genes in the other clusters. Among the four clusters that are strongly associated with only one or two genes by the chi-squared test, *TP53* is one of the five most commonly mutated genes in all four clusters (Table 2). *KRAS* is frequently mutated in three clusters; *APC* in two, and *CDKN2A* in two. It is possible that the differences between mutation clusters may be more subtle, caused by less common mutations.

Next, we turn to the clusters that are found when using Jaccard distance on the binary Newmanized transcriptomic data. Like the clusters found from the usual transcriptomic data, these clusters are well defined, have sharp boundaries, and are cleanly separated, both visually and by silhouette width (Figures 1A and 3A). One of the most interesting findings from this cluster analysis is that colon cancer (which forms part of cluster NO.1, in the lower right corner of Figures 2A and 2C, along with some cases of esophageal cancer) can be completely separated from rectal cancer (which forms part of cluster NO.2, in the lower left corner of Figures 2A and 2C, along with some cases of pancreatic cancer). Previous analyses based on the usual transcriptomic profiles have repeatedly found these two cancer cohorts to be indistinguishable [3,10,15]. (Also see the NIH web site https://www.nih.gov/news-events/nih-research-matters/colon-rectal-cancers-surprisingly-similar).

Paschke and colleagues [46] have argued that the term “colorectal cancer” should be avoided, and the two cancer types should be regarded as distinct entities. Our use of Newmanization gives strong support to their arguments, and it may provide tools to better understand the molecular differences between colon and rectal cancer. The mutation patterns, by themselves, are unlikely to fully explain the differences. Four of the five most frequently mutated genes are on both lists (Table 2). *APC* (56% vs. 49%) and *FAT4* (21% vs. 13%) mutations are more common in NO.1; *TP53* (71% vs. 61%) and *KRAS* (54% vs. 33%) mutations are more common in NO.2. At lower (but statistically significant) levels, NO.1 cases are more likely to have *BRAF* or *PI3KCA* mutations and NO.2 cases are more likely to have *NRAS* or *SMAD4* mutations.

Our gene enrichment analysis of the differences between clusters NO.1 and NO.2 were driven by pathways involving the extracellular matrix, cell adhesion, and, in particular, collagens and integrins. Rectal cancer and colon cancer have previously been reported to express different collagens, with implications for tumor progression and aggressiveness [47–49]. Rectal cancer has also been reported to have higher expression of some integrins [50]. Those results were based on direct comparisons of colon cancer and rectal cancer, using gene expression microarrays [47,50] or immunohistochemistry [48,49].

We also performed a gene enrichment analysis comparing the two NewmanOmics clusters that contained esophageal cancers. Our findings are consistent with previous reports that APOBEC mutation signatures are more common in esophageal squamous cell cancers than esophageal adenocarcinomas [51,52]. It is also known that differences in expression of HLA Class II molecules and rates of histone modifications are related to prognosis in esophageal cancer, possibly with differential effects between adenocarcinoma and squamous cell carcinoma [53–55]

### 4.1 Conclusions

Clustering using either the binary mutation data or the binary Newmanized transcriptomic data yields useful insights into the relationships between different cancers of the digestive tract. Based on our study, the strength of unsupervised clustering based on mutations is that it is able to find the set of highly mutated tumors with microsatellite instability, along with a set of tumors with very few (or no) mutations. Its weakness is that it is unable to cleanly separate the remaining sets of tumors, which have overlapping mutations in well known driver genes like *APC, TP53, KRAS, KMT2D, CDKN2A, PIK3CA*, and *SMAD4*.

The strength of unsupervised clustering using the Newmanized data is that it identifies several well-separated clusters of tumor samples with common features in the way in which their gene expression patterns differ from normal. By removing the dominant “cell-of-origin” signal from each tumor type, we were able to use an unsupervised method to separate colon cancer and rectal cancer into distinct clusters. Gene enrichment analysis using those clusters was able to recapitulate known differences that had originally been found by supervised analyses using different technologies. We were also able to separate the esophageal cancer samples into two groups that are not the usual adenocarcinoma vs. squamous cell carcinoma split, and we were able to characterize the differences in terms of differential expression of genes that could also be found in the literature.

## Supporting information

Suppplemental Table 1

Suppplemental Table 2

## 5 Acknowledgements

